# Gene Regulatory Cross Networks: Inferring Gene Level Cell-to-Cell Communications of Immune Cells

**DOI:** 10.1101/415943

**Authors:** Gokmen Altay, Bjoern Peters

## Abstract

**Background:** Gene level cell-to-cell communications are crucial part of biology as they may be potential targets of drugs and vaccines against a disease condition of interest. Yet, there are only few studies that propose algorithms on this particularly important research field.

**Results:** In this study, we first overview the current literature and define two general terms for the types of approaches in general for gene level cell-to-cell communications: Gene Regulatory Cross Networks (GRCN) and Gene Co-Expression Cross Networks (GCCN). We then propose two algorithms for each type, named as GRCNone and GCCNone. We applied them to reveal communications among 8 different immune cell types and evaluate their performances mainly via membrane protein database. Also, we show the biological relevance of the predicted cross-networks with pathway enrichment analysis. We then provide an approach that prioritize the targets by ranking them before experimental validations.

**Conclusions:** We establish two main approaches and propose algorithms for genome-wide scale gene level cell-to-cell communications between any two different cell-types. This study aims accelerating this relatively new avenue of research in cross-networks and points out the gap of it with the well-established single cell type gene networks. The proposed algorithms have the potential to reveal gene level interactions between normal and disease cell types. For instance, they might reveal the interaction of genes between tumor and normal cells, which are the potential drug-targets and thus can help finding new cures that might prevent the prevailing of tumor cells.

## Background

Gene network inference (GNI) methods can identify interactions between different genes and the gene products they encode. Gene regulatory networks (GRN), which can be inferred by GNI methods, help in the basic biological understanding of genes and their functions and are well studied within single cell condition [1]. In any GRN inference, the goal is to infer a network that consist of causal interactions between the genes. It is the main difference of GRN from the co-expression networks [2] that infers a network containing both causal and associated interactions. Given the huge number of significantly associated genes, most of the links of a co-expression network are associative. This way it can be considered mostly similar to kind of clustering of interactions among the genes in some more detail than basic clustering. In clustering, the clustered genes are assumed to be fully connected with each other. GRNs are, in theory, more precise to understand the causal mechanism of any interaction but harder to infer accurately.

However, different cell types are also interacting with each other, especially immune cell types, to manifest various complex biological functions. So far, GNI methods have not focused on the gene level interactions between different cell types [3]. In fact, it was stated in the very recent publication [3] that gene network approaches have not explicitly addressed the challenge of identifying networks representing the interaction of different cell types in a complex tissue and until then, there has been no attempt to develop a completely data-driven systems biology approach to discover novel cell-communication networks. GRNs are well studied to implement in the datasets of single cell types but, to our knowledge, we could not find a previous study that performs only GRNI, without the combination of other algorithms such as differential expression (DE), directly on the gene expression datasets of two different cell types to infer gene regulatory cross-network (GRCN) that contains causal direct interactions of genes between the two-cell types. We found a few other related studies but they are mainly the co-expression network based implementations on different cell types and those types of networks were named as *cross-talk* networks [3-5] in general. For example, in [3], a co-expression based network inference method, RELNET applied to link genes differentially expressed in two cell conditions and also introduced a gene connectivity metric to score the targets. In their application, out of 204 estimated ones, they found 107 genes (53%) are linked to the Gene Ontology term *cell communication* and therefore stated as to represent a class of proteins potentially mechanistically involved with cell cross-talk. They performed some wet-lab experiments to validate some of their findings. They also stated that the approach has the advantage to reverse engineer cell communication networks in the absence of any prior information, and in this respect, their method is different from the contemporary method of [4]. The latter has been developed to discover the interactions between stroma and cancer cells in a model of lung tumor metastases and is based on comprehensive ligand-receptor network information, which are extracted from several knowledge databases and thus requires prior information. In another algorithm [5], a comprehensive network of biomolecular interactions between human genes is compiled and used as prior information and integrated to differentially expressed genes to infer intercellular interactions between two different cell types.

The aim of our study is to establish an approach to perform blind estimation of gene-gene regulatory direct causal interactions among different cell types utilizing only the gene expression datasets of two different cell types and also show that these GRCNs provide useful and relevant biological insights. To the best of our knowledge, this is one of the first study that aims to infer causal direct regulatory interactions between genes of different cell types using only expression dataset and without the combination of any other algorithms such as DE. The closest approach might be considered as the method [3], but their algorithm is limited to infer differentially expressed (DE) genes between the two cell types. However, there are also genes that might be interacting with different genes in different cell conditions and since they express in both conditions, they may not be captured as DE genes. Our approach is solely based on dependency scores and the gene expression dataset, and thus, considers all the possible interactions on genome-wide level as described in more detail in the following.

### Gene Level Networks Between Two Different Cell Types

Conceptual approach to obtain a classical gene co-expression network (GCN) can be overviewed as follows: GCN of a single cell type is inferred by first computing an adjacency matrix among genes using a dependency estimator (e.g. Pearson Correlation Coefficient, PCC) and then eliminating statistically non-significant correlated gene pairs (links) that are below a certain dependency score threshold [2, 6, 7]. The remaining non-zero valued gene pairs are represented by a link, which make a GCN. This network can also be represented in three columns list, where first and second columns are for each gene of the pair and the third column is for the dependency score between the two genes.

A gene regulatory network (GRN) algorithm further filters the links of this GCN with its heuristic algorithm and aims to infer only direct causal interactions among the genes. For example, ARACNE breaks the weakest links of all the triplets of a GCN [8] or C3NET keeps only the maximum valued partner of each gene [9]. Establishing a similar conceptual categorization of gene networks for gene level cell-to-cell communication is also aimed in this study as we consider that might help accelerating the research in this direction. In summary, we define Gene Regulatory Cross Network (GRCN) as the causal gene level interactions between two different cell types. Whereas, we define Gene Co-expression Cross Network (GCCN) as the network that shows all the significantly co-expressed genes between the two different cell types. Additionally, the links in both GRCN and GCCN are specific to cross networks between the two cell types and cannot not appear in the individual gene networks of each of the two cell types. GCCN is similar to the widely used term cross-talk network but we make the distinction between the two as the following: GCCN only refers to gene level communications, whereas cross-talk network term is used to describe communication in any molecular levels between different cell-types. In summary, the defined GRCN and GCCN terms of this study are more specific than the well-established cross-talk networks (CTN) term that is widely used for any type of network that refers to communication between different cell types at any molecular level [10-15]. We consider that GRCN and GCCN are yet to be explored as a relatively new avenue of research and catch up with the classical gene networks inference research of single cell types. The potential of its applications is of fundamental importance. For example, it has the potential to reveal how a disease cell-type such as tumor is interacting with the normal cells around it and thus revealing the mechanism of it results new drug-targets that potentially prevent the prevailing of the tumor cells. Or, revealing gene level causal interaction mechanism among the complex immune cell types might help developing better and more specifically targeted vaccines with lowered side effects. With this motivation, we propose a new GRCN algorithm, GRCNone, and also a new GCCN algorithm, GCCNone, as described in the Methods section.

## Results

### Performance comparisons of the proposed algorithms GRCNone and GCCNone

We performed enrichment analysis of the proposed algorithms over membrane and secretome databases assuming their strong biological relevance to cell-to-cell communication. Details of the databases, datasets and the algorithms can be seen in great detail in the Methods section. Here we present their performance comparisons. We apply GRCNone over 56 different GRCN combinations of the 8 immune cell types. Therefore, in Figure 1, GRCNone box plot has 56 different Fisher’s exact test p-values. Similarly, GCCNone is applied to 28 different GCCN combinations and its box plot has 28 different p-values. We also provide the combinations of both directions for GRCNone that makes 28 different p-values too in the presented box plot to more fairly compare GRCNone with GCCNone.

**Figure 1:**
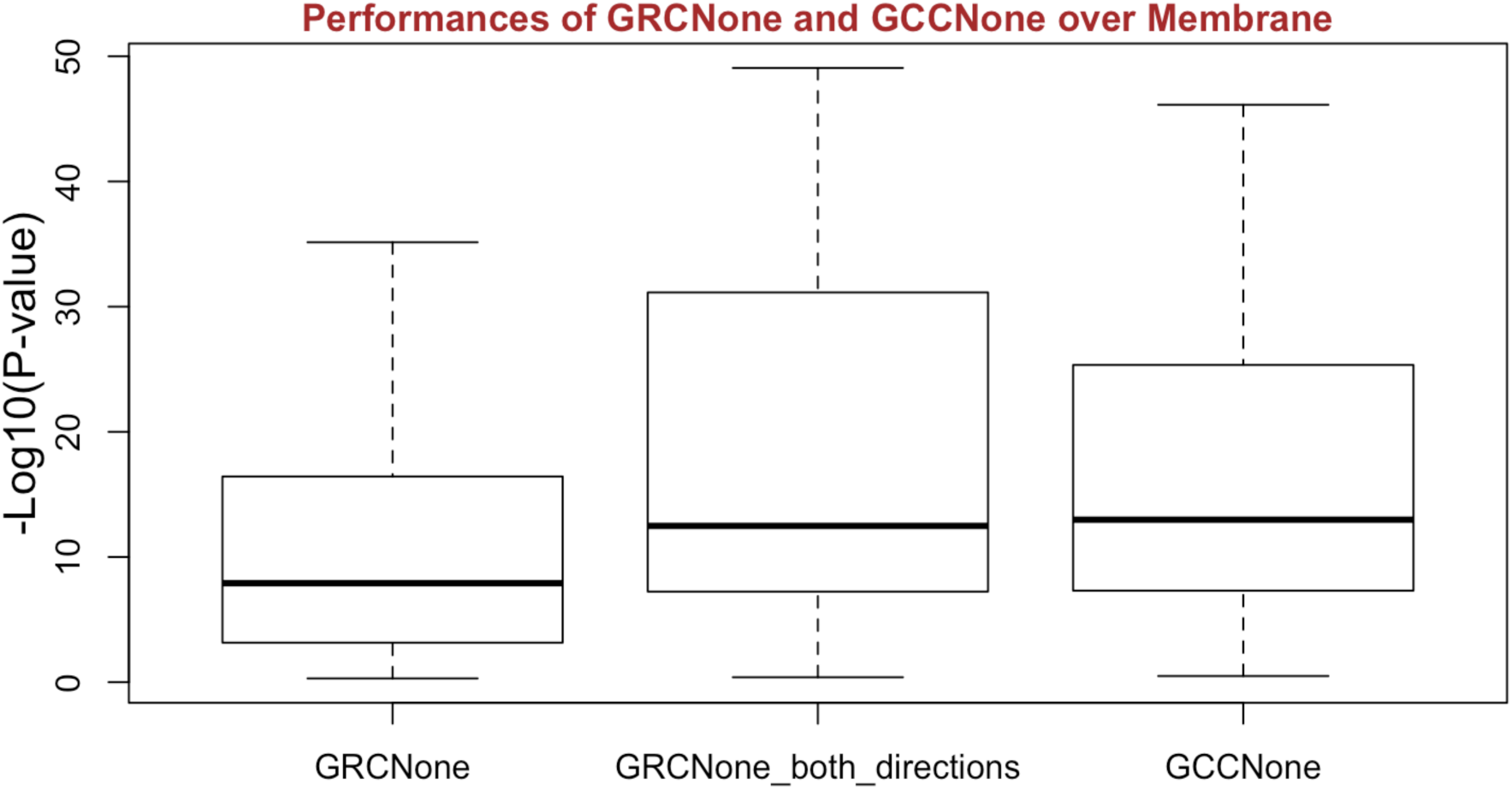
Enrichment performances of GRCNone and GCCNone algorithms along with the GRCNone both directions combined over the Membrane database. P-values were obtained by Fisher’s exact test.

The results from Figure 1 shows somewhat similar performances among the methods. We further performed statistical analysis to measure if there is any statistically significant difference. Comparing all of the three boxplots, One-way Anova test suggests that the observed difference in the means of the three different groups are statistically significant (p-value = 5.11e-03) assuming significance level, alpha, as 0.01. It is worth mentioning that GRCNone is directional and its result appears to have the lower performance. However, combinations of GRCNone network in both directions of any two different cell types (GRCNone_both_directions) and GCCNone have the same number of p-value scores (28) and it might be fairer to compare the performance of only these two results to compare GRCNone and GCCNone. Because in that case, we consider both interaction directions for both of the methods. When we perform two-sided Student’s t-test between the mean significance scores of GRCNone_both_directions and GCCNone, the p-value is 5.53e-01, (assuming alpha as 0.01) and it does not provide statistically significant difference. Thus, it suggests that there is not a statistically significant difference between both of the results of both methods regarding the Membrane database.

### GRCNone over Gene Ontology (GO) enrichment analysis

We combined all the 56 GRCNone networks among 8 cell types, which can be found in Additional file 1 and page ‘All_GRCNone’. In that, we also provided all the relevant information such as correlation values so that interested readers can filter as they wish. Using all the 5558 unique genes of both on regulator and target sides, we performed enrichment analysis over GO database (2017) of Cellular Component. The aim of this analysis was to test if the blind predictions (using only expression dataset) of GRCNone provides relevant biology. Because if it does, then it validates that the algorithm works well for the purpose as intended. If we observe unrelated biological enrichments, then it might mean that the algorithm does not work well for the purpose of cross-network inference. Since the algorithm infers GRCN, we expect to get membrane related biology. The enrichment results over GO Cellular Component (obtained using EnrichR [16]) confirmed our expectations and the validity of GRCNone algorithm. The top three of the most significant enrichments are azurophil granule membrane (GO:0035577), phagolysosome membrane (GO:0061474) and lysosomal membrane (GO:0005765) with adjusted p-values 5.61e-18, 1.66e-17, 1.02e-15 and also overlap ratios of 149/273, 141/257, 172/347, respectively. In fact, out of top 25 most significant enrichments, 15 of them contains the words with mostly *membrane* and also *transport* in their names as can be seen in Additional file 1 and page ‘All_GRCNone_GO_CC’, where we provide all the results. Therefore, enrichment analysis suggests that GRCNone provides strongly biologically relevant results for GRCN inference between two different cell types.

### GRCNone and GCCNone network sizes between the immune cells

In Figure 2, we present the number of links of each GRCNone network for all the 56 GRCN combinations of the 8 different immune cell types. From this figure, we can observe the intensity of the GRCN interactions between specific immune cell type pairs with directionality. The figure shows the intensities of interactions among the 8 immune cell types. Since the algorithm limits the number of GRCN interactions by 2000, we observe it as the maximum for only one network among the 56, which is from B-cell to Monocyte non-classical (denoted as M2). Then we observe decreasing intensity of interactions. M2 to CD8 Naive and Monocytes to CD-4 Naive and CD8 Naive in both directions have highest intensities. Lowest number of interactions are observed from CD-4 Stim to B-cell and CD-8 Stim to NK.

**Figure 2:**
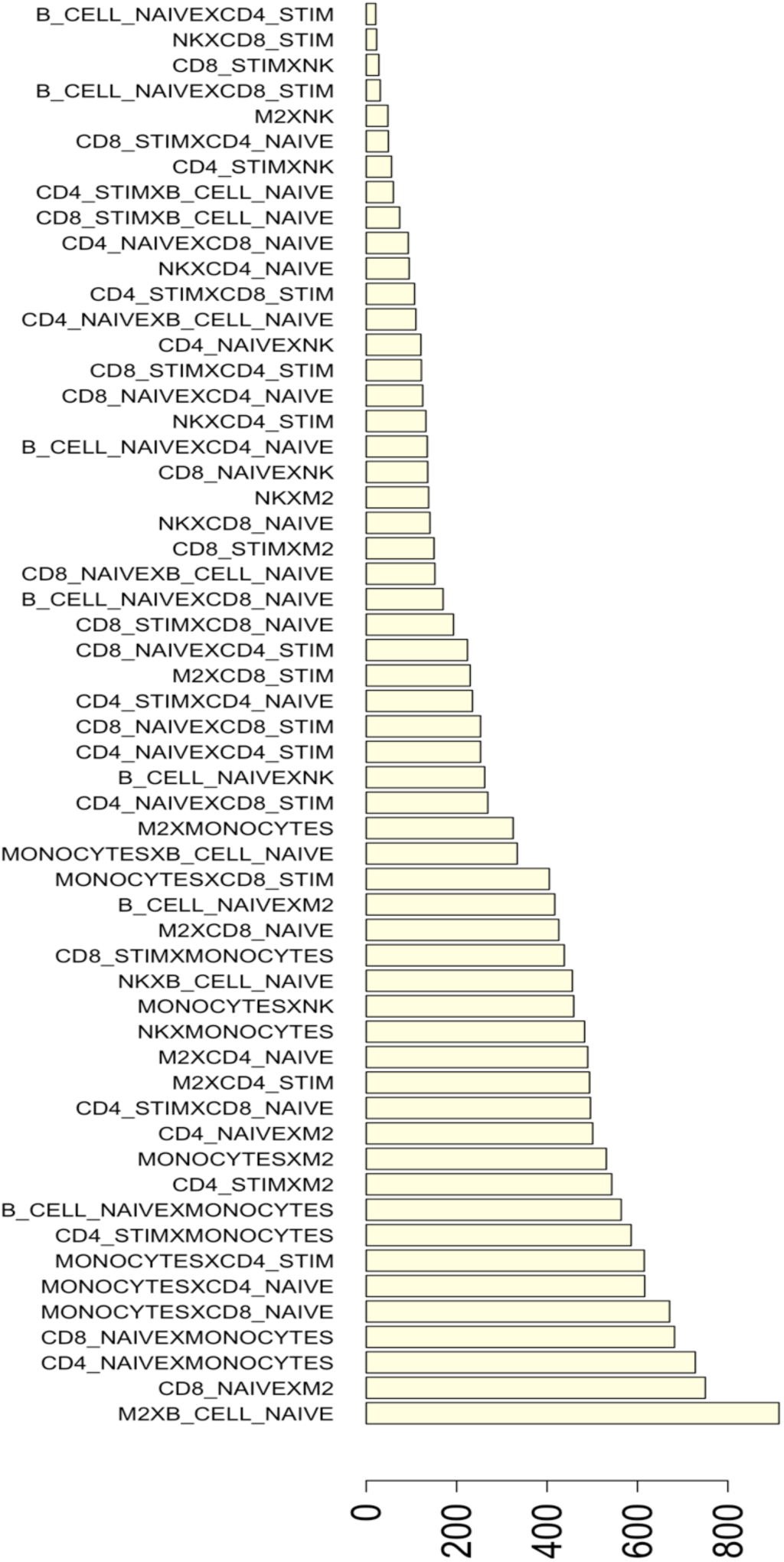
the number of links of each GRCNone networks for all the 56 GRCN combinations of the 8 different immune cell types.

In Figure 3, we present the number of links of each GCCNone network for all the 28 GCCN combinations of the 8 different immune cell types. Since the algorithm limits the number of GCCN interactions by 2000, we observe that 13 out of 28 networks have the maximum size.

**Figure 3:**
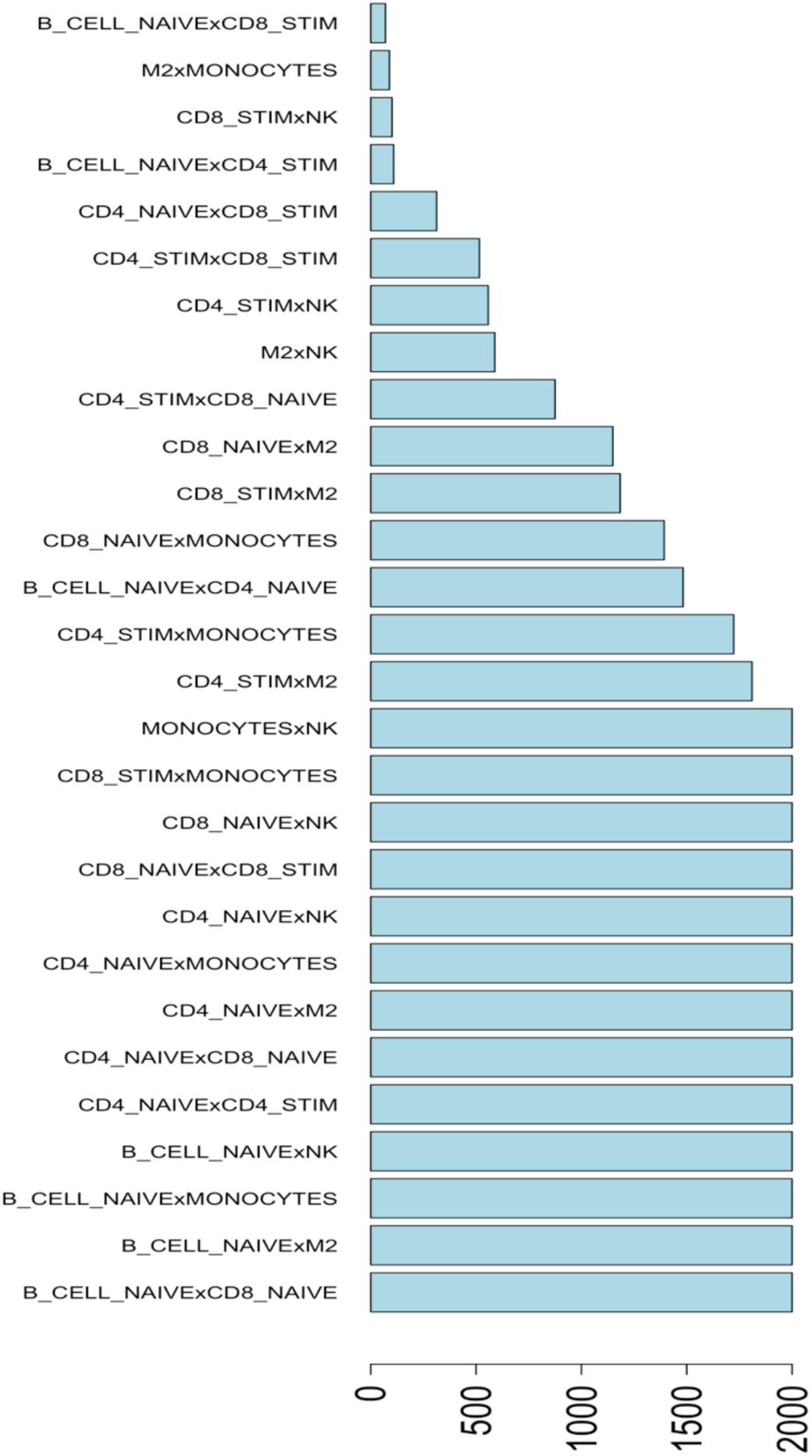
the number of links of each GCCNone networks for all the 28 GCCN combinations of the 8 different immune cell types.

As we presented previously, there is no significant difference in enrichment performances of GRCNone and GCCNone over membrane database but as it is clear that GCCNone provides higher network sizes. We selected the maximum sized network of GRCNone (M2**X**B-cell, where **X** denotes to GRCN type network) and the corresponding network of GCCNone, which is again maximum with 2000 links, and we performed general enrichment analysis to see their biological relevancies as presented in Table 1. It is worth mentioning that M2**X**B-cell GRCNone network has 913 links and1018 unique genes and it has Fisher’s exact test adjusted p-value of 1.48E-21 over the membrane database used in this study. M2xB-cell GCCNone network has 2000 links and 1185 unique genes and it has adjusted p-value of 1.45E-17 over the membrane. Since GCCNone contains both directions, in order to have a fairer comparison with GCCNone, we provide the GCRNone combined networks of M2**X**B-cell and Bcell**X**M2, which has 1316 links and 1461 unique genes and it has adjusted p-value of 7.00E-31 over the membrane database. These results suggest that GRCNone provides more membrane relevant genes than GCCNone despite the fact that it predicts lower number of interactions. When we made a general enrichment analysis over GO Cellular Component, we see from Table 1 that top ten of the results M2**X**B-cell GRCNone network are mostly membrane related whereas it is not the case in GCCNone M2xB-cell network. This observation suggests that GRCNone provides biologically more specific relevant results.

**Table 1:** Enrichments analysis (via EnrichR over GO Cellular Component) of GRCNone network from M2 to B-cell types and GCCNone network between M2 and B-cell.

| GRCNone |  |  | GCCnone |  |  |
| --- | --- | --- | --- | --- | --- |
| **G O C e l l u l a r C o m p o n e n t** | Adj. P-value | Overlap | **G O C e l l u l a r C o m p o n e n t** | Adj. P-value | Overlap |
| phagolysosome membrane (GO:0061474) | 0.000794 | 32/257 | nuclear speck (GO:0016607) | 3.63423E-12 | 80/525 |
| lysosomal proton-transporting V-type ATPase complex (GO:0046611) | 0.000794 | 29/226 | nucleolar part (GO:0044452) | 1.15816E-06 | 75/635 |
| cytolytic granule membrane (GO:0101004) | 0.000794 | 29/226 | nucleolus (GO:0005730) | 1.15816E-06 | 75/637 |
| azurophil granule membrane (GO:0035577) | 0.000794 | 32/273 | paraspeckles (GO:0042382) | 1.15816E-06 | 41/260 |
| endolysosome membrane (GO:0036020) | 0.000794 | 29/233 | Cajal body (GO:0015030) | 1.15816E-06 | 43/281 |
| lysosomal HOPS complex (GO:1902501) | 0.000794 | 28/222 | omega speckle (GO:0035062) | 1.15816E-06 | 43/278 |
| extrinsic component of lysosome membrane (GO:0032419) | 0.000794 | 28/222 | Gemini of coiled bodies (GO:0097504) | 2.48954E-06 | 40/259 |
| integral component of lysosomal membrane (GO:1905103) | 0.000794 | 28/223 | LYSP100-associated nuclear domain (GO:0016606) | 3.24583E-06 | 39/256 |
| **nucleolar part (GO:0044452)** | 0.0099179 | 54/635 | **PML body (GO:0016605)** | 3.92722E-06 | 43/302 |
| cytoplasmic side of lysosomal membrane (GO:0098574) | 0.0008926 | 28/226 | sphere organelle (GO:0071601) | 3.24583E-06 | 39/256 |

### Literature search of some the top predicted targets

#### Top regulator hubs

We combined all the regulator hubs (with >= 3 targets) of the 56 GRCNs among the 8 immune cell types and ranked them based on the number of targets regulated by them, which can be seen in Additional file 1 at page “Regultors_ALL”. We performed literature search to see if the top 5 regulators have any related literature.

The top regulator hub is RP11-425L10.1 pseudogene that regulates 639 targets from B CELL NAIVE to non-classical monocyte (M2). In a very recent study [17], RP11-425L10.1 was shown to be differentially expressed when comparing untreated lymphocytes of BRCA1/2 mutation carriers and controls. It was among the 81 upregulated genes of which the enrichment analysis resulted significant enrichment in the T-cell receptor signaling, immune system, and B-cell receptor signaling pathways. We accept this literature evidence as a strong support for the prediction as it shows its involvement in B-cell receptor signaling pathway and also as a pseudogene, it would yet to be explored further in the future.

Second top regulator hub is MIR4426 pseudogene that regulates 322 targets from B CELL NAIVE to NK cells. In [18], MIR4426 was shown to be among the microRNAs targeting TLR (Toll-like receptors) pathway over TLR8. Toll-like receptors are a specialized group of membrane receptors that detect pathogen-associated conserved structures and also dysregulation in the TLR signaling can possibly result in a dysregulated immune response which could contribute to major intestinal pathologies including colorectal cancer [18]. This reference shows the evidence of the role of MIR4426 immune response and thus stands as a support for the prediction.

Third top regulator hub is TNFRSF1A protein coding gene that regulates 169 targets from M2 to B CELL NAIVE cells. It is also in both membrane and secretome databases and these evidences are sufficient to support the prediction but there is also countless literature for its role in immune system. Forth top regulator hub is SUSD3 protein coding gene that regulates 158 targets from CD4_NAIVE to M2 cells is also in membrane databases. Similarly, fifth top regulator hub is C14orf2 protein coding gene that regulates 127 targets from CD8-Stim to M2 cells is also in membrane databases. For all the top 5 targets, we found evidence from the literature that supports the predictions of GRCNone algorithm. We leave the exploration of the rest of the hubs to the interested readers.

It is worth mentioning that, the above top 5 regulator hub targets are the blind predictions of GRCNone algorithm; namely with no prior information at all and based solely on the expression datasets. However, since the membrane and secretome databases are available, one can further filter the results to find more reliable experimental targets as will be discussed in the coming sub-sections. This might be especially helpful for selecting better experimental targets from the predicted hubs with small number of targets (e.g. <10).

### Some of Top Experimental Targets

The top hub regulators as discusses in the previous subsections are primary experimental targets in general for wet-lab validation experiments. However, since wet-lab experiments are expensive to conduct, we prefer to further filter the results by selecting interactions with the pairs of genes that are in membrane or secretome databases as well. We combined all the 56 GRCNone networks and kept links that correspond the above criterion and also ranked them by the number of remaining targets of hubs. There are 752 such links left, which can be seen in Additional file 1 on page “Targets_SecMemBoth”.

The largest hub is TNFRSF1A with 45 interactions of which 42 of them from M2 to B-cell. IL6ST is the top experimental target of TNFRSF1A with highest correlation magnitude (−0.72). IL6ST protein functions as a part of the cytokine receptor complex (ncbi.nlm.nih.gov/gene/3572) and mediates signals which regulate immune response (uniprot.org/uniprot/P40189). Mutation in IL6ST causes immunodeficiency and a pleiotropic defect in signaling by multiple cytokines (including IL-6, IL-11, IL-27, and OSM) and also associated with defective B cell memory [19]. The literature somewhat validates our prediction, given the strong support of both IL6ST and TNFRSF1A in immune cells, thus they appear to be promising experimental targets for inter-cellular interaction between M2 and B-cells.

We selected the first experimental target of TNFRSF1A but it has 44 more targets and it would be helpful to prioritize the targets before experimenting. We already ranked them by correlations and this might help if there was not any related prior information. However, in order to utilize from the literature, we performed enrichment analysis via EnrichR [16] of the 44 targets that are in B-cell. We expected to observe a related biology to accept as validation. B Cell Receptor Signaling Pathway_Homo sapiens_WP23 appeared (WikiPathways) as the top enriched pathway (p-value= 0.017) with the two target genes CD79B and LAT2. Therefore, these two are selected as the next promising experimental targets of TNFRSF1A among the others.

The second top hub regulator in the list is SUSD3 with 32 interactions of which 42 of them from CD4-Naive to M2 cells. SUSD3 has already a known role in cell-cell interactions (genecards.org) and there is a patent on it (US20110038875A1) showing its expression on the surface of CD4 T cells. NFAM1 is the top experimental target of SUSD3 with the highest correlation magnitude (0.7) CD4-Naive to M2 cells. The protein encoded by NFAM1 is a type I membrane receptor that activates cytokine gene promoters such as the IL-13 and TNF-alpha promoters (ncbi.nlm.nih.gov/gene/150372). Innate immune system is among the related pathways of NFAM1 and Gene Ontology (GO) annotations related to it include transmembrane signaling receptor activity (genecards.org). Given the strong support from the literature, regulation of NFAM1 by SUSD3 from CD4-Naive to M2 cells is selected as another important experimental target. There are 31 other targets of SUSD3, which are left to the attention of interested readers. We performed enrichment analysis of the 32 targets (in M2 cells) of SUSD3 and in Jensen Tissues database, monocyte appeared as the top enriched tissue type of the set (p-value = 1.034e-7). Also, TOR Signaling_Homo sapiens_WP1471 appeared (WikiPathways) as the top enriched pathway (p-value= 0.0015) with the two target genes TSC2 and RPTOR, which can be the other experimental targets of SUSD3.

There is also another interesting regulator, GIMAP5 with 13 interactions, mostly from NK cells to B-cells or CD8-Naive cells. It is interesting because one of its direct target is IFNGR1 and stands as an easier experimental target. Also, GIMAP5 is already a known immune related gene. For example, Gimap5 deficient mice showed reduced IL-7rα expression, and T-dependent and T-independent B cell responses are abrogated [20].

In this section, we have manually selected some of the most promising experimental targets for further wet-lab validation studies of the predictions of our proposed algorithm GRCNone. We leave all the other targets to the attention of the interested readers in the Additional file 1 on page “Targets_SecMemBoth”, with which a similar experimental target selection approach can be performed.

### Discussions and Conclusion

We define methodologies, GRCN and GCCN, for gene level cell-to-cell communication on the genome-wide scale and propose two algorithms, GRCNone and GCCNone, which corresponds to each methodology. We implemented the algorithms on the 8 different immune cell types. The results have significant validation by the literature. We foresee that this study and the proposed methods will accelerate the progress of this relatively new avenue of research field on the genome-wide scale gene level communications between different cell types. As a consequence, it will help revealing novel biological insights between cell-to-cell interactions on the gene level in general. The potential and variety of the applications of the proposed methods are very large. Therefore, we expect significant contributions of the current study to biology in general. For instance, it might help revealing the causal gene level interaction from a disease cell-type to the normal cell type around it, which might cause exploring new drug-targets that might prevent the prevailing of the tumor cells. In the case of immune cell-types, revealing genome-wide scale gene level causal interaction mechanism among the complex immune cell types might help developing better targeted vaccines with lower side effects.

This is an ongoing study and we plan to extend it by performing wet-lab experiments for the validations of some of the important targets found in this study.

## Methods

In this section, we describe all the methods and datasets and databases used for the analysis of this study.

### RNA-seq Datasets of Immune Cells

We first describe the datasets used in the application of the proposed algorithms. We used RNA-seq datasets of 8 different immune cell types. This data was produced by the DICE (Database of Immune Cell Expression, Expression quantitative trait loci (eQTLs) and Epigenomics) project (dice-database.org). Its GEO accession number and citation will soon be updated in the DICE database. DICE project produced RNA-Seq data for 8 different immunological cell types with ∼100 samples each, ∼15 of which were repeat samples gathered from the same donor (Table 2). The cell types are CD4 Naive T cell (CD3+ CD4+ CD45RA+ CD127+ CCR7+), CD4 Stim T cell (stimulation with aCD3/CD28 for 4 hours after FACS sorting), CD8 Naive T cell (CD3+ CD8+ CD45RA+ CD127+ CCR7+), CD-8 Stim T cell (stimulation with aCD3/CD28 for 4 hours after FACS sorting), Monocyte classical (CD14+ CD16-), Monocyte non-classical (CD14low CD16+), NK cells (CD3-CD56dim CD16+) and B cell naive (CD19+ CD20+ CD24low CD38low CD27-IgD+ IgM+ IgG-)

**Table 2:**
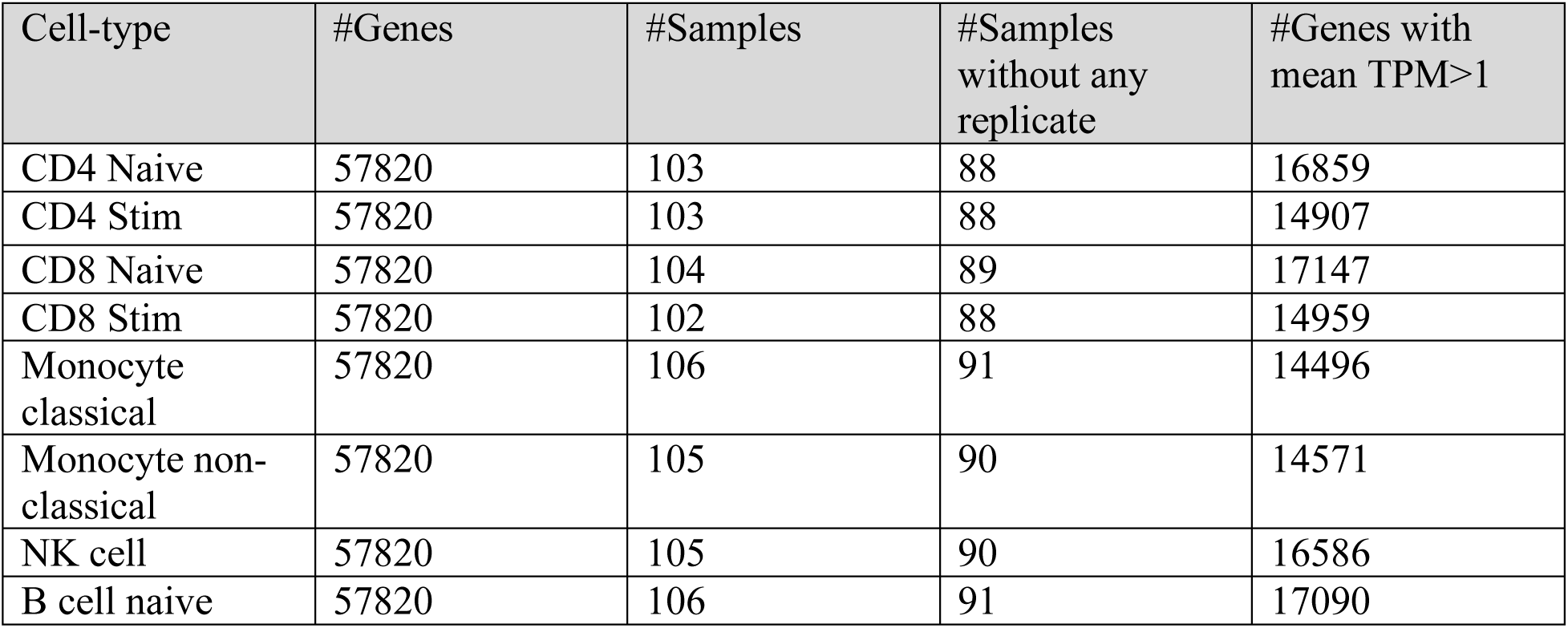
The number of genes and samples of all the 8 cell types used for the analysis.

The workflow that generated the RNA-seq count data from raw files are summarized as follows: For each RNA-Seq dataset, the single-end reads that passed Illumina filters were filtered for reads aligning to tRNA, rRNA, adapter sequences, and spike-in controls. The reads were then aligned to UCSC mm9 reference genome using TopHat (v 1.4.1) [21]. DUST scores were calculated with PRINSEQ Lite (v 0.20.3) [22] and low-complexity reads (DUST > 4) were removed from the BAM files. The alignment results were parsed via the SAMtools [23] to generate SAM files. Read counts to each genomic feature were obtained with the htseq-count program (v 0.6.0) [24] using the “union” option. After removing absent features (zero counts in all samples), the raw counts were used for bioinformatics analysis. Sample sizes of the datasets might be considered as sufficient for GNI analysis [25].

### The Reference Database for Performance Evaluation: The human secretome and membrane proteome databases

For the performance evaluations, we need a reference database to evaluate the predictions of the algorithms. We used the human secretome and membrane proteome database as it is biologically most relevant to cell-to-cell communications. The following information and the databases were obtained from http://www.proteinatlas.org/humanproteome/secretome by June 2017 [26]. A membrane protein is associated or attached to the membrane of a cell or an organelle inside the cell and can be classified as either peripheral or integral. A secretory protein can be defined as a protein which is actively transported out of the cell. 39% of the 19628 human protein-coding genes are predicted to have either a signal peptide and/or at least one transmembrane region suggesting active transport of the corresponding protein out of the cell (secretion) or location in one of the numerous membrane systems in the cell. Several genes code for multiple protein isoforms (splice variants) with alternative locations, including 675 genes with both secreted and membrane-bound isoforms (downloaded and referred as sm database in our manuscript). 2918 genes (15%) are predicted to have at least one secreted protein product (downloaded and named as s database). Also, 5455 genes (28%) are predicted to have at least one membrane-bound protein product (downloaded and referred as m database in our manuscript). A large fraction of the clinically approved treatment regimes today use drugs directed towards (or consisting of) secreted proteins or cell surface-associated membrane proteins. Since s, m and sm databases are biologically important and more relevant to inter-cellular communications, we used them as reference in our enrichment analysis to evaluate the performance of our GRCN predictions. Secretome list, s, has 2918 proteins but 1333 of them are in the universe set of our data. Membrane list, m, has 5455 proteins but 2909 of them are in the universe set of our data. Intersection of Secretome and membrane list has 675 proteins but 389 of them are in the universe set of our data. We used Fisher’s exact test for the overlapping analysis. Part of the membrane proteins are named as transmembrane proteins (TP), which is a type of integral membrane protein that spans the entirety of the biological membrane to which it is permanently attached. Many transmembrane proteins function as gateways to permit the transport of specific substances across the biological membrane [26].

### Gene Regulatory Cross Network Algorithm: GRCNone

We aimed to develop a general, unbiased from pre-knowledge, GRCN algorithm that provides genome-wide scale causal directional interactions between the genes of two different cell types using only gene expression datasets. Since it is the first attempt specifically for the defined GRCN inference, we named the proposed algorithm as GRCNone.

In brief, with a single interaction example between the arbitrarily named genes GeneA5 of cell A and GeneB3 of cell B, GRCNone infers a directional causal interaction from GeneA5 of cell A to GeneB3 of cell B if the correlation value between the two genes is statistically significant and GeneA5 is the maximum absolute correlation (magnitude) valued partner of GeneB3. This criterion is searched for each gene of cell B with respect to all the genes of cell A between the two cell types and an initial raw GRCN from cell type A to cell type B is obtained. Then the raw GRCN network is compared with the two individual GCNs of the two cell types and only the significantly different ones are kept. The final result is a GRCNone network that also shows the direction of the cross-network from cell A towards cell B. A detailed illustration is shown in Figure 4 along with the following in depth explanation.

**Figure 4:**
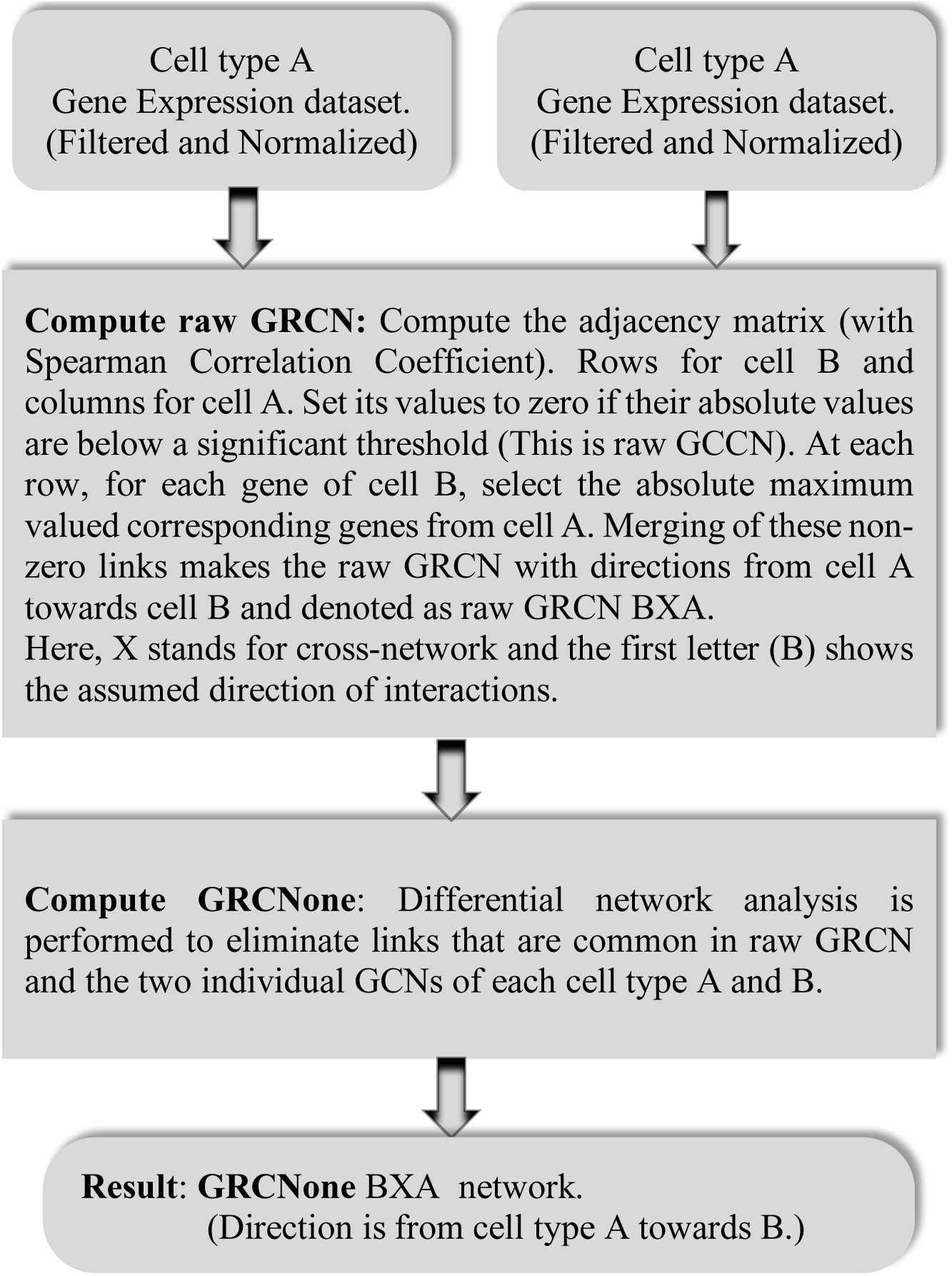
General block diagram of the introduced GRCNone algorithm.

Figure 4 provides an overview of the proposed GRCNone algorithm. Here, we will explain it in more detail. Two gene expression datasets of two different cell types are used as input. The goal is to find the specific genes of cell-type A that directly regulates or causes a direct effect on the specific genes in cell-type B during the communication between the two cell-types. This means, we assume that there is a biologically cell-to-cell communication from cell-type A towards B. Let’s say cell-type A and B have the expression datasets sizes of M rows (genes) with S columns (samples) and N rows with S columns, respectively. M and N are the number of genes in cell-type A and B, respectively. Independent from any high-level analysis, such as network inference of this study, data is optionally preprocessed to reduce the effect uninformative noisy variables. Everyone can choose their own way of preprocessing the data before inputting into the high-level algorithms. After the filtering preprocess step, in general each dataset is left with different number of genes. In our analysis, for filtering, we kept genes with at least one average TPM (Transcript Per Million) across all the samples in each dataset. Therefore, the sizes of N and M are in general different than each other. S is the sample size of both of the datasets with the same sample order. We also normalized the datasets separately with the Variance-stabilizing transformation (VST) [27] of the popular DeSeq2 R package [28]. As we said, this preprocessing step is just suggested and fully optional and can be changed or not performed if wished. These preprocessed datasets are input to ‘Compute raw GRCN’ step as seen in Figure 4. We then compute a rectangular adjacency matrix of size N rows and M columns by calculating the Spearman Correlation Coefficients (SCC) between the genes of both of the cell-types. In the end of this section, we provide a more detailed analysis result that caused the selection of the SCC and VST combination in GRCNone. These components are optional in using GRCNone and they can be exchanged in the future if better alternatives are explored by the users. The proposed algorithm, GRCNone, is meant to be the general approach but not the particular components used in it. After obtaining the rectangular adjacency matrix, we then filter out the weak correlation values by setting them to zero in the adjacency matrix if the absolute of SCC correlation value is below 0.6 that is selected as a loose threshold for general analysis. Alternatively, if computational cost is not an issue, one could obtain another threshold via an arbitrarily selected p-value based on the null distribution that can be generated by permutation analysis similarly as described in [7]. Since the final results are all ranked, the user can increase the threshold to more stringent values after the initial analysis and focus on smaller number of stronger targets. We also provide the effect of various thresholds on performances in the end of this section that helped us select SCC threshold as 0.6 in our dataset.

Then, the non-zero values in the adjacency matrix represent the co-expressed gene pairs between the two cell types. Until this step, all the processes are the same as with our second proposed algorithm GCCNone that will be described in the following sub-section. In order to not repeating the explanations so far, we named the attained adjacency matrix at this point as ‘raw GCCN’ adjacency matrix so that we can refer to it and continue while describing GCCNone in the following sub-section.

Afterwards, we select the maximum correlated partners (genes from cell type A) for each gene on the rows (genes of cell type B) in the raw GCCN adjacency matrix. Computationally, this is implemented by taking the maximum valued column (or columns if there are multiple maximums) for each row in the raw GCCN adjacency matrix of size NxM. This is inspired by the proposed hypothesis that a gene is most likely to be regulated by its maximum correlated partner gene [29]. The resulting network is called ‘raw GRCN’ that includes the links from cell type A towards cell type B at the gene level. For the given example cell types, the raw GRCN network can be denoted as raw GRCN B**X**A, where ***X*** here stands for the cross-network that is a keyword for cell-to-cell communication. In this denotation, the direction of the links is from cell type A towards cell type B. The raw in the name represents that the network is not yet filtered for the links that are also available in each of the co-expression networks of the individual cell types. This raw GRCN network is then input to the ‘Compute GRCNone’ step as seen in Figure 4.

At this step, rank based differential network analysis is performed to eliminate the links of the raw GRCN that might also appear similarly in both of the individual gene co-expression networks of each cell type. In the raw GRCN BXA, for each gene of the B side, we rank the correlations of its corresponding partner genes in the adjacency matrix between the two cell-types. The highest absolute correlation is ranked as 1. Therefore, in the raw GRCN BXA, the rank of all the genes at the B side with their counterparts at the A side is 1 as they are already the maximum absolute correlation values. We also compute the rank of all the genes at the A side with their counterparts at the B side in the same adjacency matrix between the two cell types but in this case the ranks are computed for each column in the matrix because the genes of cell type A are on the columns. This provides two columns ranks list that show the ranks of the correlation magnitudes of each gene pair with respect to each other in the raw GRCN BXA. For the same gene pairs of the raw GRCN, we then generate the ranks of them in the adjacency matrices of the individual cell types A and B if the same gene pairs are available. Then, we compare the rank differences between the cross-network and individual ones. We set the rank difference to be at least 60 (Additional File 1 contains performances of for various rank differences and correlation thresholds). If both of the ranks of the raw GRCN gene pairs have difference above 60 with compare to their ranks in both of the individual cell types then they are kept as GRCN specific gene pairs, namely links, with the following additional condition that must hold at the same time: If the magnitude of the cross link correlation value is at least 10% more than the magnitude of the individual within cell correlations (assuming they have the same sign), then we select that cross link as they also satisfy the rank criterion. This assures that the rank difference also represents some level of correlation magnitude value increase. We also have an additional independent criterion that selects all the links based on only their magnitudes as this may represent some extreme cases, such as positive and negative correlation values of the same link (cross and within cell) but with similar magnitudes. That is, if the magnitude of the correlation values difference, between cross-link and the link of a single cell type, is greater or equal than the maximum of the either of the correlation magnitudes, then that link is also selected regardless of the rank criteria. We apply it for all the interactions in the raw GRCN BXA.

All the resulting selected links compose of the final regulatory cross-network, which is GRCNone BXA, where the links are considered to be from cell type A towards B. This direction assumption is hypothetical and might be re-considered upon our follow-up experimental validation studies. As a final rule, we further limit the maximum number of interactions with 2000 and thus make sure that we do not infer extreme number of cross-network interactions but only most important portion.

As an application side note, assuming that the interactions of raw GRCN are directional from A to B, half of the rank comparisons, means for only B side, could be enough but the assumption of the direction is hypothetical. Therefore, in order to make sure we only have GRCN specific links, we compare all the cross-network ranks with all the corresponding ranks in adjacency matrices of both of the individual cell-types.

We provide performance results for various rank differences and correlation thresholds values in Additional File 1 for both GRCNone and GCCNone. It is worth mentioning that, raw GRCN parts are inspired from Ac3net [29] and C3NET [9] of single cell type GRNI algorithms. Moreover, differential analysis part is also inspired from DC3NET [30]. GRCNone is based on modification, adaption and integration of those well-established single cell type GRNI algorithms for GRCN inference.

### Gene Coexpression Cross Network Algorithm: GCCNone

We developed a general, unbiased from pre-knowledge, GCCN algorithm that provides cross-network specific co-expression network between the genes of two different cell types using only gene expression datasets. Since it is one the first attempt specifically for the defined GCCN inference, we named the proposed algorithm as GCCNone that is inspired and developed based on RELNET [2] and DC3NET [30] by modification, adaption and integration of those well-established single cell type GNI algorithms for GCCN inference.

GCCNone and GRCNone has common steps until the point where ‘raw GCCN’ adjacency matrix is obtained as explained in the previous section where GRCNone is explained in detail. However, the difference of GCCNone from GRCNone is that after that step GCCNone does not perform the rest of the process of the second step (‘Compute raw GRCN’ block as illustrated in Figure 4) and just passes to the third step that is the same as GRCNone differential network analysis. The process of GCCNone algorithm is illustrated in Figure 5 below for completeness. The resulting GCCNone network between cell types A and B can be denoted as GCCNone BxA or GCCNone AxB, which means there is no directionality. The lower case x is to emphasize that this cross-network is a co-expression cross-network, namely GCCN.

**Figure 5:**
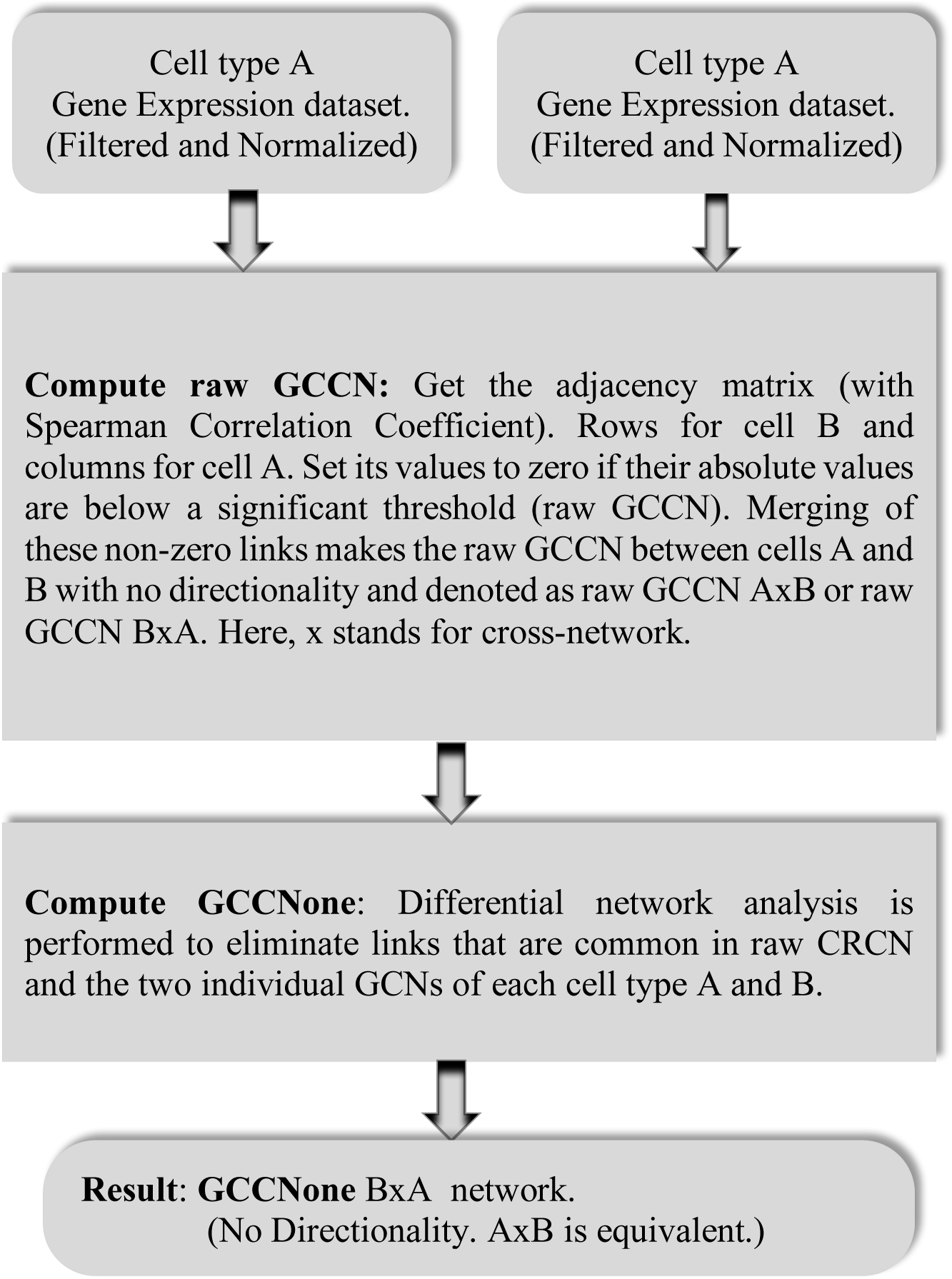
General block diagram of the introduced GCCNone algorithm.

### Preprocessing Combinations Selection for GRCNone

We implemented and analyzed the performance of some prominent preprocessing components to select the optimum combination in order to use them in GRCNone. For the dependency scores estimator, we implemented the popular estimators the Pearson Correlation Coefficient (PCC) and the Spearman Correlation Coefficient (SCC). A comprehensive review and descriptions of dependency score estimators can be seen in [31]. Shortly, PCC captures linear relationship between the variables, whereas SCC is a ranked based version of PCC and captures monotonic relationships. As the sign of the dependency scores is considered as important, we did not prefer mutual information based estimators.

For the data normalizations, we first implemented TMM (the trimmed mean of M-values normalization) [32], which is the normalization function of the popular edgeR [33] R package. We also implemented Variance-stabilizing transformation (VST) [27] of the popular DeSeq2 R package [28], and lastly the simple Logarithm of base 2. They already made 6 possible combinations and their performances can be seen in Figure 6.

**Figure 6:**
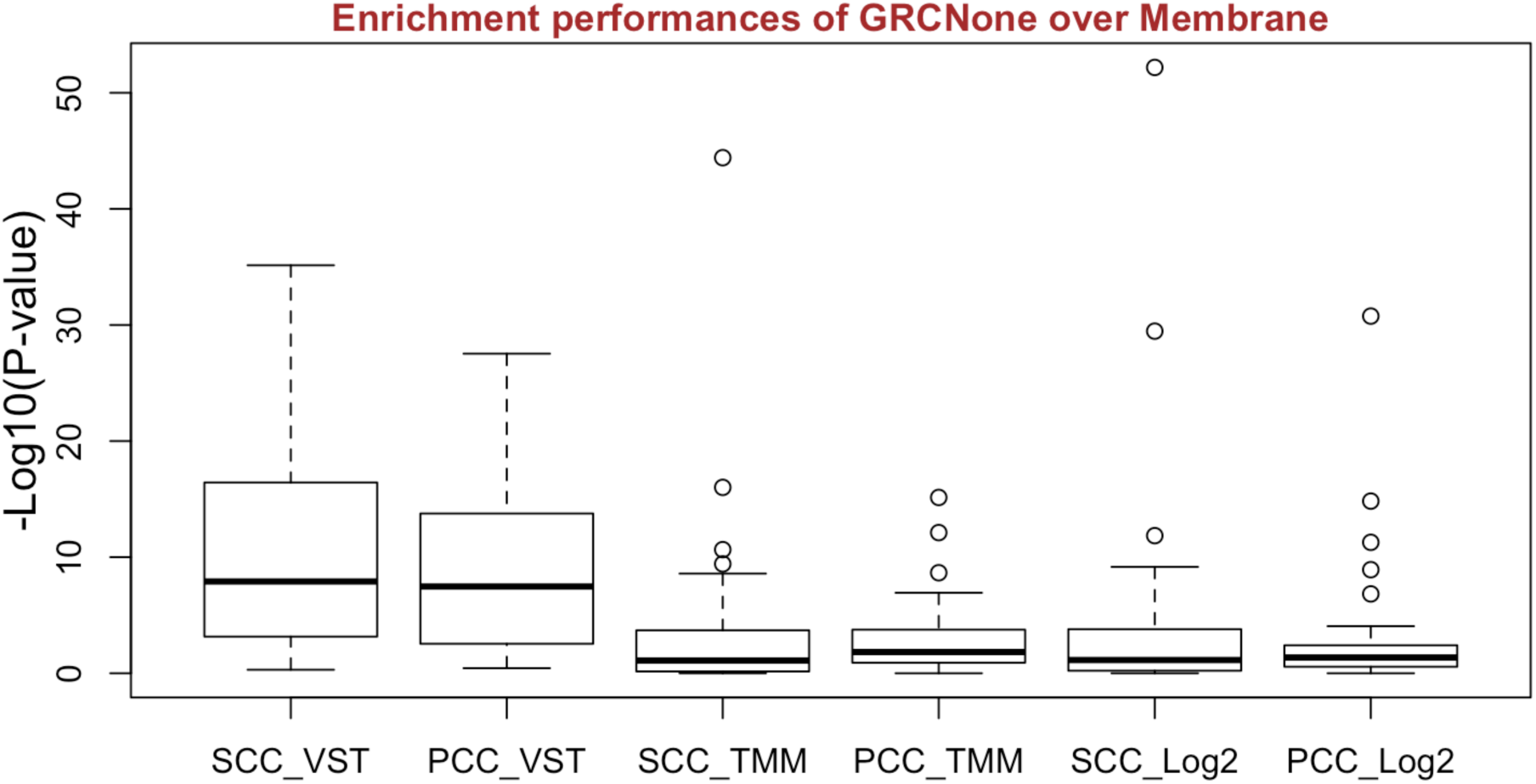
Enrichment performances of GRCNone over membrane with all the 6 possible preprocessing combinations.

SCC and VST combination has the highest median and maximum scores. One-way Anova test suggests that the observed difference in the means of the different groups are statistically significant (p-value = 2.75e-13). However, when we compare the difference with two-sided t-test on the means of SCC_VST and PCC_VST, then the p-value is 3.31e-01. It means that the difference between SCC_VST and PCC_VST is not statistically significant (considering alpha = 0.01). We selected SCC as the estimator as it is less affected from the variations of the data and a default choice as a correlation estimator in most of the real-life applications. In fact, as shown below we also perform the similar comparison analysis for GCCNone algorithm, where SCC_VST combination again provides the best median and best maximum performance values and there is also statistically significant difference among the combinations. As a summary, based on the preprocessing combinations performances seen in Figure 6, we select the SCC as the estimator and VST as the normalization method for the preprocessing of the RNA-seq data before inputting into our GRCNone cross regulatory network inference algorithm.

### Preprocessing Combinations Selection for GCCNone

We also implemented and analyzed the performance of some preprocessing components to select optimum combination for GCCNone as shown in Figure 7. We apply GCCNone over 28 different cross co-expression network combinations of the 8 cell types. Therefore, in Figure 7, each box plot has 28 different p-values.

**Figure 7:**
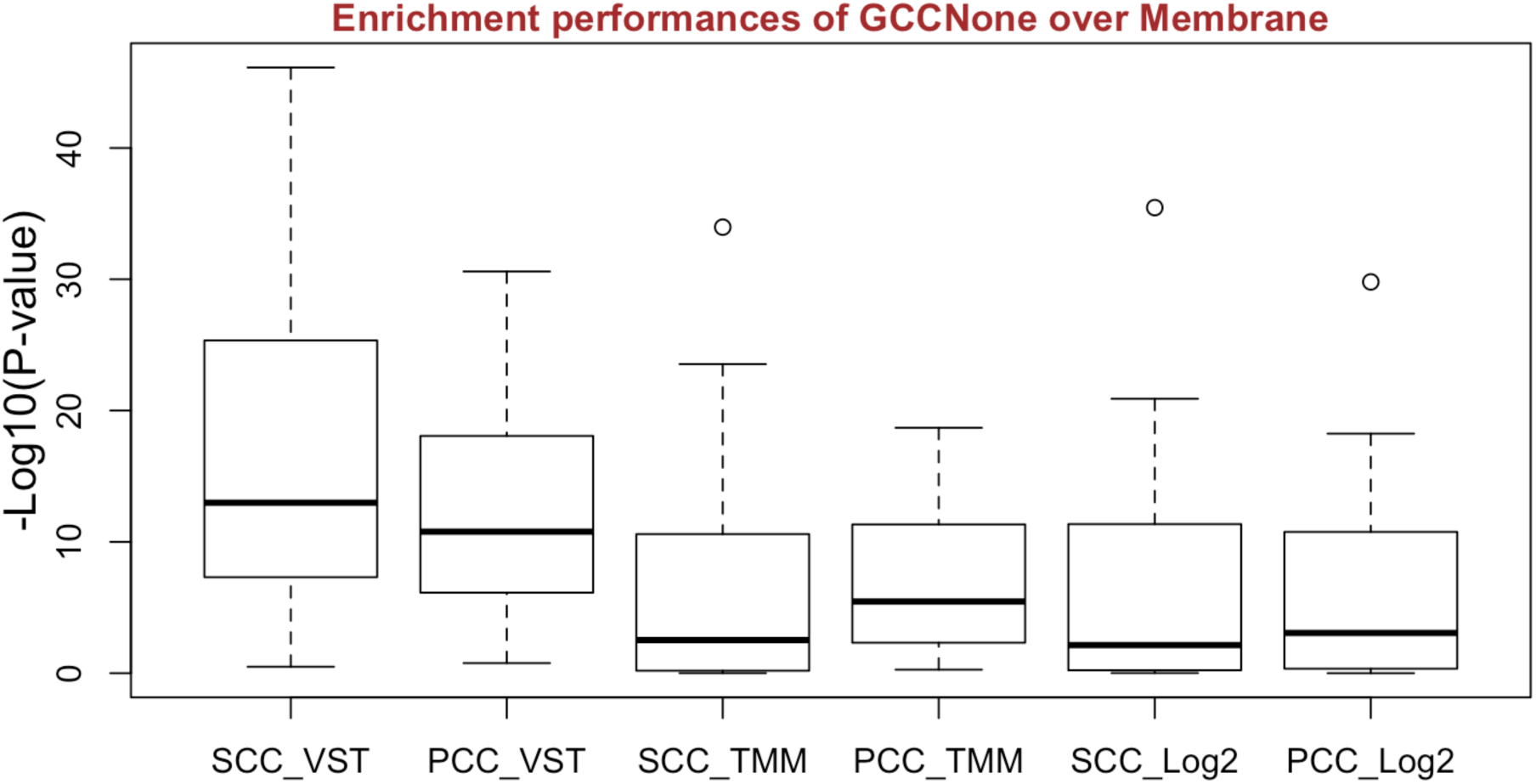
Enrichment performances of GRCNone over membrane with all the 6 possible preprocessing combinations.

One-way Anova test suggests that the observed differences in the means of the different groups are statistically significant (p-value = 2.33e-05). When we compare the difference with two-sided t-test on the means of SCC_VST and PCC_VST, then the p-value is 2.10e-01. It means that the difference between SCC_VST and PCC_VST is not statistically significant (considering alpha = 0.05). However, SCC_VST combination provides best median and best maximum performance values. This support the decision to select SCC and VST combination in GCCNone algorithm too. As a summary, based on the preprocessing combinations performances seen in Figure 7, we select the SCC as the estimator and VST as the normalization method for the preprocessing of the RNA-seq data before inputting into our GRCNone cross regulatory network inference algorithm and GCCNone cross co-expression network inference algorithm.

### List of abbreviations

BS: B-spline
CPM: counts per million
CS: Chao-Shen
CT: copula transformation
GCCN: Gene Co-Expression Cross Networks
GNI: Gene network inference
GRCN: Gene Regulatory Cross Networks
GRN: Gene regulatory networks
GSEA: Gene Set Enrichment Analysis ()
Log-2 or l2: Logarithm with base 2
noCT: no copula transformation
NoNorm or none: No normalization (also denoted as *nonnormalized*)
QN or Q: quantile normalization (QN)
PCC: Pearson Correlation Coefficient
SCC: Spearman Correlation Coefficient
PBG: Pearson-based Gaussian
RLE: relative log expression (RLE)
TMM: The trimmed mean of M-values normalization
TPM: transcript per million
VST: Variance-stabilizing transformation

**#** symbol stands for “number of”

### Declarations

The content is solely the responsibility of the authors and does not necessarily represent the official views of the National Institutes of Health.

## Acknowledgements

We used R [34] for the programming. We thank Brendan Ha and Jason Greenbaum for providing the information about the used datasets of DICE project in this study and help writing in the subsection ‘RNA-seq Datasets of Immune Cells’.

## Funding

Research reported in this publication was supported by National Institute of Allergy and Infectious Diseases (NIAID) of the National Institutes of Health (NIH) under award number: R24AI108564.

## Authors’ contributions

BP conceived and coordinated the study, contributed to the algorithm design, analysis and writing up the manuscript. GA designed the GRCNone and GCCNone algorithms and performed all the analysis, developed and run the software programs and wrote the manuscript. All authors read and approved the final manuscript.

## Competing interests

The authors declare that they have no competing interests.

